# Deep Learning Based Segmentation of Brain Tissue from Diffusion MRI

**DOI:** 10.1101/2020.07.30.228809

**Authors:** Fan Zhang, Anna Breger, Kang Ik Kevin Cho, Lipeng Ning, Carl-Fredrik Westin, Lauren J. O’Donnell, Ofer Pasternak

## Abstract

Segmentation of brain tissue types from diffusion MRI (dMRI) is an important task, required for quantification of brain microstructure and for improving tractography. Current dMRI segmentation is mostly based on anatomical MRI (e.g., T1- and T2-weighted) segmentation that is registered to the dMRI space. However, such inter-modality registration is challenging due to more image distortions and lower image resolution in the dMRI data as compared with the anatomical MRI data. In this study, we present a deep learning method that learns tissue segmentation from high-quality imaging datasets from the Human Connectome Project (HCP), where registration of anatomical data to dMRI is more precise. The method is then able to predict a tissue segmentation directly from new dMRI data, including data collected with a different acquisition protocol, without requiring anatomical data and inter-modality registration. We train a convolutional neural network (CNN) to learn a tissue segmentation model using a novel augmented target loss function designed to improve accuracy in regions of tissue boundary. To further improve accuracy, our method adds diffusion kurtosis imaging (DKI) parameters that characterize non-Gaussian water molecule diffusion to the conventional diffusion tensor imaging parameters. The DKI parameters are calculated from the recently proposed mean-kurtosis-curve method that corrects implausible DKI parameter values and provides additional features that discriminate between tissue types. We demonstrate high tissue segmentation accuracy on HCP data, and also when applying the HCP-trained model on dMRI data from a clinical acquisition with lower resolution and fewer gradient directions.

## 1. Introduction

Segmentation of brain tissue types, e.g., the labeling of gray matter (GM), white matter (WM), and cerebrospinal fluid (CSF), is a critical step in many diffusion MRI (dMRI) visualization and quantification tasks. Most current tissue segmentation approaches are based on T1-weighted (T1w) or T2-weighted (T2w) anatomical MRI data (Ashburner and Friston, 2005; Fischl, 2012; Smith et al., 2004), which has high image resolution and image contrast that differentiates between tissue types. However, application of anatomical-MRI-based segmentation to dMRI requires inter-modality registration, which is challenging since dMRI often has echo-planar imaging (EPI) distortions (Wu et al., 2008; Albi et al., 2018; Jones and Cercignani, 2010) and low image resolution (Malinsky et al., 2013). Improved dMRI acquisitions can mitigate these challenges. For instance, the Human Connectome Project (HCP) acquired high-resolution dMRI data with alternating phase encoding to correct EPI distortions, making it easier to register between the anatomical MRI data and the dMRI data (Glasser et al., 2013). However, advanced HCP-like acquisitions are not always available and are not yet feasible for many clinical settings where scan time is limited. Therefore, there is a need for segmentation approaches that can be applied on the dMRI data directly. Such approaches would be useful for MRI protocols that are missing high quality anatomical data, or when it is difficult to achieve high quality registration of the anatomical-MRI-based segmentation to the dMRI space.

Most dMRI-based brain tissue segmentation methods use features derived from the diffusion tensor imaging (DTI) model, such as mean diffusivity (MD) and fractional anisotropy (FA) (Liu et al., 2007; Schnell et al., 2009; Wen et al., 2013; Kumazawa et al., 2013; Yap et al., 2015; Ciritsis et al., 2018; Zhang et al., 2015; Nie et al., 2018). These DTI-based studies have achieved limited accuracy partly because DTI is known to be non-specific in characterization of water diffusion in the complex intracellular and extracellular in vivo environment (O’Donnell and Pasternak, 2015). Diffusion kurtosis imaging (DKI) (Jensen and Helpern, 2010), a clinically feasible extension of DTI that characterizes non-Gaussian water molecule diffusion, enhances DTI by providing information about molecular restrictions and tissue heterogeneity in the brain. DKI parametrizes the dMRI data with a diffusion tensor and a kurtosis tensor (Tabesh et al., 2011). From the kurtosis tensor, additional parameters are derived, including mean kurtosis (MK), axial kurtosis (AK) and radial kurtosis (RK), that can quantify the complexity of the microstructural environment of different brain tissues (Steven et al., 2014).

Segmentation methods that include DKI parameters show improved brain tissue segmentation compared to DTI (Beejesh et al., 2019; Hui et al., 2015; Steven et al., 2014). However, one important challenge is that DKI parameters can be affected by signal alterations caused by imaging artifacts such as noise, motion and Gibbs ringing (Veraart et al., 2016; Shaw and Jensen, 2017; Zhang et al., 2019). Consequently, DKI often yields output parameter values that are implausible (e.g. see MK in Fig. 1(a2)), which affects DKI-based analyses including tissue segmentation (Veraart et al., 2016; Shaw and Jensen, 2017). However, correction methods such as the mean-kurtosis-curve (MK-Curve) method (Zhang et al., 2019) are now available to robustly identify and correct implausible kurtosis tensor parameter values (Zhang et al., 2019).

**Figure 1:**
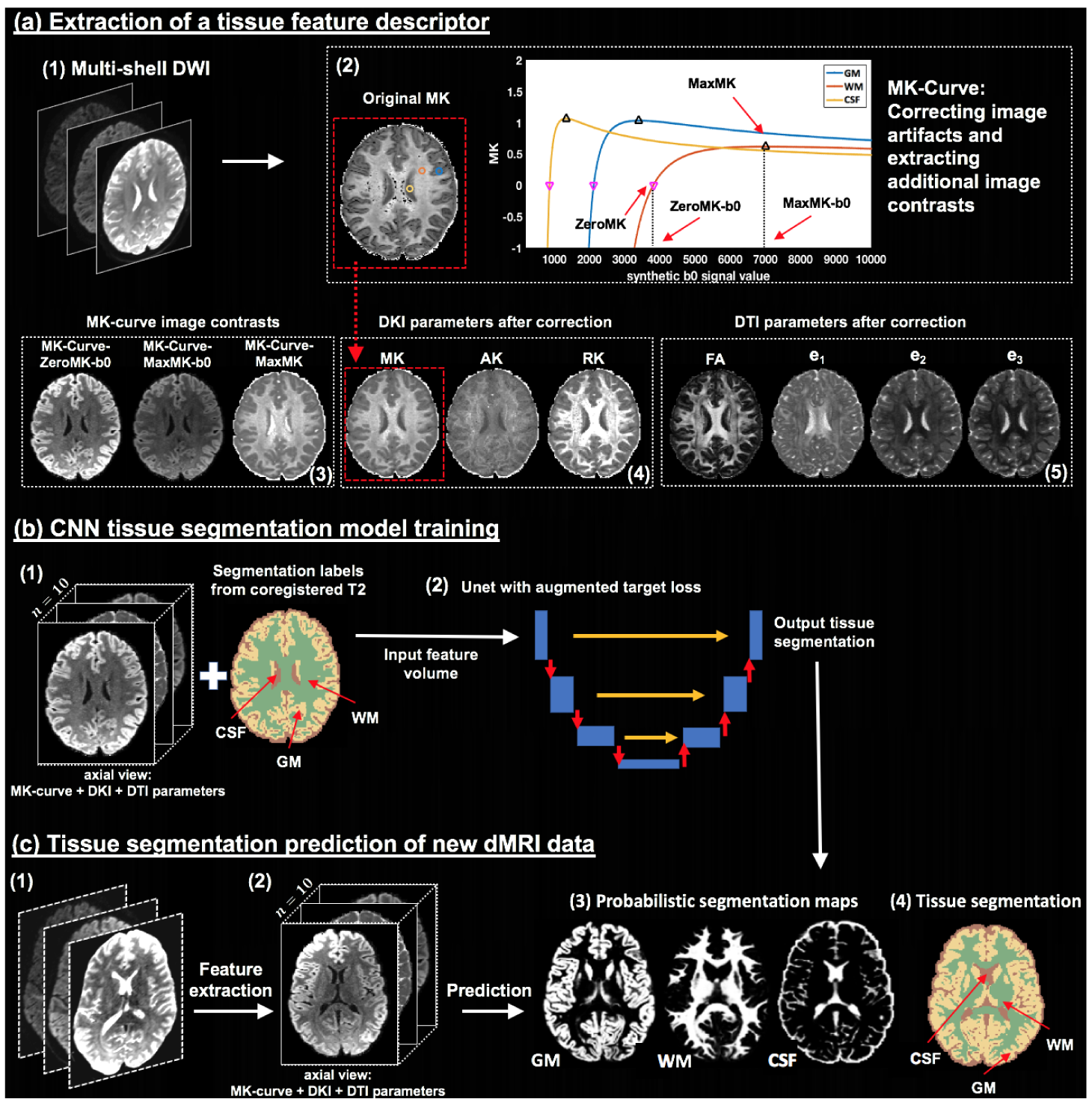
Method overview. Given input DWI data (a1), an MK-Curve is computed for each voxel (a2, showing three example voxels). The MK-Curve is used to correct implausible DKI and DTI parameters, and to derive three additional image contrasts that are useful to identify tissue types: ZeroMK-b0, MaxMK-b0 and MaxMK (a3). A total of 10 features are computed: the 3 MK-Curve contrasts (a3), 3 DKI maps (a4) and 4 DTI maps (a5). The corrected images no longer have implausible values (e.g., original MK map (a2) versus corrected MK map (a4); in a red frame). The 3D volumes of the 10 features along with the segmentation labels computed from co-registered T2w data (b1) are used to train a CNN with a Unet architecture and with a novel augmented target loss function (b2). For new DWI data (c1), the trained CNN is applied on the computed 3D volume of the 10 features in the dMRI space directly (c2), not requiring inter-modality registration. The final prediction output includes a probabilistic segmentation map for each tissue type (c3) and a segmentation label per voxel (c4).

Recent advances in deep learning methods have shown significant improvement for solving image segmentation problems (e.g., see reviews in Akkus et al. (2017); Bernal et al. (2019); Garcia-Garcia et al. (2017). The most widely used deep learning model is the convolutional neural network (CNN) that is designed to automatically and adaptively learn spatial hierarchies of image features from low-to high-level patterns (LeCun et al., 1998; Krizhevsky et al., 2012). So far CNN for brain segmentation have been considered as part of multi-modal segmentation that includes dMRI, showing highly promising tissue segmentation performance (Zhang et al., 2015; Nie et al., 2018). However, this sort of use still highly depends on the accuracy of cross-modality registration that is required to form the multi-modal input used for the segmentation.

In this paper, we propose a deep learning approach that utilizes a CNN to learn the segmentation of WM, GM, and CSF directly from dMRI features while obviating the need for inter-modality registration. The CNN is trained using high quality and coregistered anatomical and dMRI data (from the HCP), and then predicts tissue segmentation of new subjects directly from dMRI data, without the need for anatomical MRI data nor inter-modality registration. To further improve accuracy we trained the CNN using a novel augmented target loss function (Breger et al., 2020) that penalizes segmentation errors in tissue boundary regions. The dMRI input includes seven DKI and DTI parameter maps that have been corrected for implausible values using MK-Curve and three additional MK-Curve-derived maps (Zhang et al., 2019). We trained, validated and tested the proposed method using 100 high-quality and co-registered anatomical and dMRI datasets from HCP, and then demonstrated generalizability to dMRI data from a clinical acquisition with lower resolution and fewer gradient directions. Overall, quantitative and qualitative comparisons with several state-of-the-art segmentation methods show that our method provides highly accurate and reliable dMRI tissue segmentation performance.

## 2. Methods

The proposed method includes 3 main steps (overviewed in Fig. 1): (a) extracting DKI- and MK-Curve-based tissue feature descriptor (Section 2.1), (b) training a CNN model for tissue segmentation (Section 2.2), and (c) predicting subject-specific tissue segmentation from new dMRI data (Section 2.3).

### 2.1. Extraction of a tissue feature descriptor

Feature extraction from the DWI data was performed using a DKI model fit (Tabesh et al., 2011), in combination with generating the MK-Curve for each voxel (Zhang et al., 2019), to correct implausible DKI and DTI parameter values, and to provide three additional image contrasts that are useful for tissue segmentation.

MK-Curve is a continuous plot that shows the dependence of MK on variation in the b0 signal (Zhang et al., 2019). In brief, an MK-Curve is generated for each voxel by replacing the original b0 value with a range of synthetic values, and calculating MK (Tabesh et al., 2011) for each signal realization. The resulting curve is characterized by three features for each voxel (Fig. 1(a2)): *ZeroMK-b0* – the largest b0 value where MK crosses 0; *MaxMK-b0* – a b0 value larger than ZeroMK-b0 where MK reaches maximum; and *MaxMK* – the MK value at MaxMK-b0. These three features define ranges of b0 values that generate implausible MK values. Implausible values can then be corrected by projecting out-of-range b0 values to the plausible range (Zhang et al., 2019), and refitting the DKI model (Tabesh et al., 2011) using the projected b0 value. This process results with corrected DKI and DTI maps that no longer have physically implausible values and fewer visually implausible voxels (e.g., comparing dark voxels in the original MK in Fig. 1(a2) with the corrected MK in Fig. 1(a4); in a red frame).

As a result of this process, for each subject, we calculate a tissue feature descriptor that is a 4D image with 10 features per voxel (Figs. 1(a3 to a5)): 3 MK-Curve features (ZeroMK-b0, MaxMK-b0, and MaxMK), 3 MK-Curve-corrected DKI parameters (MK, AK and RK) and 4 MK-Curve-corrected DTI parameters (FA, and the three eigenvalues, e1, e2, e3). Each parameter was rescaled using a z-transform across all voxels within a brain mask.

For comparison (see Section 3.2.1) we also considered the following descriptors: 1) a DKI-based descriptor that included only the DKI and DTI features obtained by fitting the DKI model to the original, uncorrected data. 2) a DTI-based descriptor, which was computed by nonlinear fit of the b=0 and b=1000 s/mm^2^ data to a diffusion tensor.

### 2.2. CNN tissue segmentation model training

We applied CNNs to train models that segment WM, GM and CSF from different sets of dMRI input parameters. In brief, the CNN had the typical three layers (LeCun et al., 1998; Krizhevsky et al., 2012): a convolution layer and a pooling layer to perform feature extraction, and a fully connected layer to map the extracted features into the final tissue segmentation maps. There were two main CNN design choices: 1) choice of CNN architecture, and 2) design of loss function. These two choices are described below.

#### 2.2.1 Multi-view 2D Unet CNNs

For the CNN architecture, we selected a Unet architecture (Ronneberger et al., 2015), which has been successfully applied for neuroimage analysis (Dong et al., 2017; Wasserthal et al., 2018; Hwang et al., 2019). While in principle the Unet architecture allows extensions to 3D image segmentation, we used 2D image slices as network input to boost memory efficiency and to enable processing of the high-resolution HCP data used for model training (see Section 3.1 for data details). However, to leverage the 3D neighborhood information in the 3D feature volume, we trained three 2D CNNs using 2D slices from three different orientations: axial, coronal and sagittal. This is similar to the model training process proposed in Wasserthal et al. (2018). The final tissue segmentation prediction was computed based on the average of the prediction probabilities across the three views (See Supplementary Material 1 for a comparison using the three views and using each individual view).

#### 2.2.2. Augmented target loss function

A loss function, which evaluates how well the CNN models the training data, is needed to optimize the weights of the CNN. To further improve the CNN training, we designed a new loss function according to the recently proposed framework of augmented target loss functions (Breger et al., 2020). Augmented target loss functions modify a given loss function by including prior knowledge of a particular task. The prior knowledge is introduced by using transformations to project the prediction (output *y*) and the ground truth (target *t*) into alternative spaces (Breger et al., 2020). In its general form, the augmented target loss function, *L*_*AT*_, can be written as

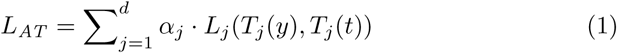

where *T*_*j*_ is a transformation, *L*_*j*_ is a loss function, and *α*_*j*_ *>* 0 is a weight for each of the *d* loss functions *j* ∈ {*1,…, d*}.

To improve the segmentation of GM, WM and CSF, we add two kinds of prior information: 1) misclassified voxels are more common on the boundaries of the different regions, and 2) CSF has essentially different diffusion properties (free diffusion) from GM and WM (hindered and restricted diffusion). Therefore, we added two penalty elements to the CNN’s loss function:

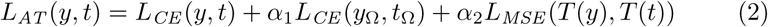

Here, the first term, *L*_*CE*_, is the conventional categorical cross-entropy loss function (Goodfellow et al., 2016). The second term penalizes misclassification in voxels included in the boundary region, Ω, which was computed by applying a Laplacian of Gaussian (LoG) edge filter (Parker, 2010) to the target segmentation, *t*. The third term penalizes misclassification between CSF and either GM or WM by applying the transformation *T* (**·**) = ⟨**·**, (1, 1, 0)^*T*^ ⟩ on *y* and *t*, where ⟨·, ·⟩ denotes the inner product and the three channels correspond to WM, GM and CSF, respectively. *L*_*MSE*_ is the mean-squared error loss function, i.e., (*T* (*y*) *- T* (*t*))^2^ over all samples. The added information/penalization is weighted by the parameters *α*_1_ and *α*_2_.

We trained three Unet CNN models, corresponding to the axial, coronal and sagittal views of the input volume, using the Deep Learning Toolbox^*T M*^ in Matlab (version 2018). Adaptive moment estimation (ADAM) was used for optimization, with a learning rate of 0.0005, a batch size of 8, and a total of 50 epochs.

### 2.3. Tissue segmentation prediction of new dMRI data

To perform tissue segmentation on new subjects that were not included in the training, we applied the trained CNN model to the dMRI data of these subjects (Fig. 1(c)). First, tissue feature descriptor was extracted from the new dMRI data (as described in Section 2.1), resulting in a 4D volume of the 10 features (Fig. 1(c2)). Zero padding was performed to maintain the same matrix size as the training images. Then, the 4D volume was separated to axial, sagittal and coronal slices, and each slice was fed to the trained CNN model of the corresponding view to output segmentation probability maps with a probability score for each tissue type. For each voxel, these scores were averaged across the output of the three views. These prediction outputs were then re-constructed into 4D maps (Fig. 1(c3)). From the 4D probabilistic segmentation maps, we computed a 3D tissue segmentation map (Fig. 1(c4)), where each voxel was assigned with the tissue type that had maximal probability.

## 3. Experiments

### 3.1. Datasets

We included two kinds of MRI datasets. The first kind was high-quality MRI data from the HCP (Glasser et al., 2013). This data was used to train the CNNs, as well as for model validation and testing. The second kind was MRI data with a clinical acquisition protocol (CAP). The CAP data was used to test how the trained CNNs generalized to data from a different acquisition. For each dataset, we used both dMRI data and anatomical T2w data. The dMRI data was used to extract the MK-Curve-based feature descriptor. The T2w data was used to generate tissue segmentation that will be considered as “ground truth”. As mentioned above, the high quality of the HCP data allows accurate registration of the T2w-based segmentation to the dMRI space, resulting with a high quality segmentation in the dMRI space that can be considered close to ground truth. For the CAP datasets, the co-registered T2w-based segmentation may not be optimal, but we use it here as a comparison to maintain a consistent reference point. Below, we introduce the MRI acquisition, data preprocessing, and T2w-based tissue segmentation for each dataset.

#### 3.1.1. Human Connectome Project (HCP) dataset

We included MRI data from a total of 100 HCP subjects (age: 29.1 *±* 3.7 years; gender: 54 females and 46 males), where 70 subjects were assigned for model training, 20 subjects were assigned for validation, and 10 subjects were assigned for testing.

The HCP MRI data was acquired with a high quality image acquisition protocol using a customized Connectome Siemens Skyra scanner. The acquisition parameters used for the dMRI data were TE = 89.5 ms, TR = 5520 ms, phase partial Fourier = 6/8, and voxel size = 1.25 *×* 1.25 *×* 1.25 mm^3^. A total of 288 images were acquired for each subject (acquired in both LR and RL phase encoding to correct for EPI distortions), including 18 baseline images with a low diffusion weighting b = 5 s/mm^2^ and 270 diffusion weighted images evenly distributed at three shells of b = 1000/2000/3000 s/mm^2^. The acquisition parameters for the T2w data were TE = 565 ms, TR = 3200 ms, and voxel size = 0.7 *×* 0.7 *×* 0.7 mm^3^. The dMRI data has been processed following the well-designed HCP minimum processing pipeline (Glasser et al., 2013), which includes brain masking, motion correction, eddy current correction, EPI distortion correction, and coregistation with the anatomical T2w data.

To obtain input tissue type labels for training and testing, we applied Statistical Parametric Mapping (SPM, version SPM12) (Ashburner and Friston, 2005) on the T2w images, resulting in a CSF, WM or GM label for each voxel. The labels were projected into the co-registered dMRI data using nearest neighbor interpolation. We note that SPM is often the method of choice in generating high quality reference brain segmentation to compare with dMRI-based tissue segmentation (Schnell et al., 2009; Ciritsis et al., 2018; Cheng et al., 2020), although other segmentation tools, e.g., FMRIB’s Automated Segmentation Tool (FSL FAST) (Jenkinson et al., 2012) could also be used instead.

#### 3.1.2. Clinical Acquisition Protocol (CAP) dataset

The CAP dataset contained MRI data from 10 healthy subjects (age: 22.7 *±* 5.4 years; gender: 5 females and 5 males) for testing the training CNN tissue segmentation model. This dataset was acquired with approval of the local ethics board.

The CAP MRI data was acquired using a Siemens Verio 3T scanner (Magnetom Verio; Siemens Healthcare, Erlangen, Germany). The acquisition parameters used for the dMRI data were TE = 109 ms, TR = 15800 ms, phase partial Fourier = 6/8, and voxel size = 2 *×* 2 *×* 2 mm^3^. A total of 125 images were acquired for each subject, including 5 baseline images with b = 0 s/mm^2^ and 60 diffusion weighted images evenly distributed at two shells of b = 1000/3000 s/mm^2^. The acquisition parameters for the T2w data were TE = 422ms, TR = 3200 ms, and voxel size = 1 *×* 1 *×* 1 mm^3^ using a 3D T2-SPACE sequence The dMRI data was processed using our in-house data processing pipeline, including brain masking using the SlicerDMRI extension (dmri.slicer.org) (Norton et al., 2017; Zhang et al., 2020) in 3D Slicer (www.slicer.org), eddy current-induced distortion correction and motion correction using FSL (Jenkinson et al., 2012), and EPI distortion correction and coregistation with the anatomical T2w data using the Advanced Normalization Tools (ANTS) (Avants et al., 2009). Manual quality check was performed to confirm good registration performance.

The T2w-based tissue segmentation of the CAP dMRI data was computed in the same way as the HCP data. SPM12 was used to compute the T2w-based tissue labels, which were projected to the co-registered diffusion space using a nearest neighbor interpolation sampling.

### 3.2. Experimental comparison

We evaluated the performance of our proposed CNN based segmentation against: 1) other machine learning classifiers, and when using different dMRI feature descriptors, and 2) three state-of-the-art dMRI-based tissue segmentation methods.

#### 3.2.1. Comparison of tissue feature descriptors and machine learning classifiers

We compared 9 approaches that used different tissue feature descriptors and machine learning classifiers. The compared feature descriptors were: 1) *F*_*DTI*_-containing the 4 DTI parameters computed from DTI modeling (as in (Kumazawa et al., 2010)); 2) *F*_*OrigDKI*_ - containing the 3 DKI and 4 DTI parameters from DKI modeling (without any correction for implausible parameter values) (as in (Beejesh et al., 2019)); and 3) *F*_*Proposed*_ - containing all 10 MK-Curve-based parameters. For each of these feature descriptors, we performed tissue segmentation using three different classifiers: 1) *SVM* - a support vector machine classifier with a radial basis function (RBF) kernel function, as used in Ciritsis et al. (2018); Schnell et al. (2009), 2) *CNN* - a Unet architecture CNN with the conventional categorical cross-entropy loss function, 3) *CNN* _*AT*_ - a Unet architecture CNN with our augmented target loss function. For each compared classifier, hyperparameters were well-tuned following a cross-validation process on the validation dataset. Specifically, in the proposed *CNN* _*AT*_, the weighting parameters *α*_1_ and *α*_2_ were set to 0.05 and 0.5, respectively, as determined during the cross-validation. In total, there were 9 comparison methods in this experiment.

For each of the compared methods, we computed a tissue segmentation model using the 70 HCP datasets assigned for training, where the hyperparameters of the model were tuned using the 20 HCP datasets assigned for validation. To evaluate the tissue segmentation prediction performance, we applied each trained model on the 10 HCP and the 10 CAP datasets designated for testing, and we calculated the segmentation prediction accuracy, *ACC*, against the T2w-based labels for each dataset, as:

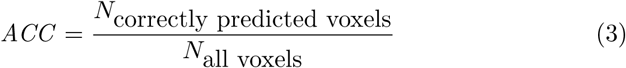

We computed *ACC* across voxels from the entire brain, and also separately across voxels from the tissue boundary and from non-boundary voxels (detected using a LoG filter). For each of the regions (entire brain, boundary and nonboundary), the mean and the standard deviation of the segmentation prediction accuracy across the 10 HCP and the 10 CAP datasets, respectively, were computed for each compared method.

#### 3.2.2. Comparison with other state-of-the-art dMRI tissue segmentation methods

We evaluated the proposed method with comparison to three other state-of-the-art dMRI-based tissue segmentation methods: 1) segmentation based on the estimation of a dMRI-based tissue response function (Dhollander et al., 2016, 2018) in the constrained spherical deconvolution (CSD) framework (Tournier et al., 2007), 2) application of the SPM tissue segmentation algorithm (Ashburner and Friston, 2005) directly to the diffusion baseline (b0) image, 3) a method that uses the direction-averaged diffusion weighted imaging (daDWI) signal as input to SPM tissue segmentation (Cheng et al., 2020). In the rest of the paper, we refer to these methods as the *CSD* method, the *SPM+b0* method, and the *daDWI* method, respectively. Briefly, the CSD method is an unsupervised algorithm that leverages the relative diffusion properties to estimate response functions^1^ of GM, WM and CSF (Dhollander et al., 2016). Subsequently, the response functions are used in the multi-shell multi-tissue CSD (MSMT-CSD) algorithm (Tournier et al., 2007) to calculate volume fraction maps for the three tissue types (Jeurissen et al., 2014), which are analogous to the probabilistic maps that our proposed method outputs. The SPM+b0 method leverages the fact that the b0 has a T2w contrast and thus applies the SPM tissue segmentation method (Ashburner and Friston, 2005) directly on the b0 image without inter-MR-modulity registration. The SPM+b0 method was previously used, for example, to assist brain tissue segmentation (Mah et al., 2014). The daDWI method fits a power-law based model to the direction-averaged signal of the DWI images to obtain two parametric maps named alpha and beta (McKinnon et al., 2017). The alpha and beta images are used for creating a pseudo T1w image (Cheng et al., 2020), which is subsequently inputted to SPM to obtain the GM, WM and CSF segmentation. For each of he compared methods, the output of the prediction includes the overall tissue segmentation map where each voxel has a predicted label (GM, WM, CSF or background) and three probabilistic segmentation maps (for GM, WM and CSF, respectively) where each voxel has a probability belonging to a certain tissue type.

We evaluated our proposed method against the CSD, the SPM+b0, and the daDWI methods by comparing the overall *ACC*. We also assessed spatial overlap with the co-registered T2w-based “ground truth” segmentation for each individual tissue type, *T* =(WM, GM, CSF), by computing the Dice score (Dice 1945) of the predicted segmentation, *Seg* _*PD*_, and the ground truth segmentation, *Seg* _*GT*_, as:

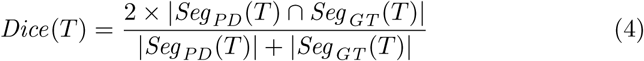

where |*Seg* _*PD*_ *∩ Seg* _*GT*_ |indicates the number of voxels in the intersection of the predicted and the ground truth segmentations, and |*Seg* _*PD*_ |and |*Seg* _*GT*_ |indicate the number of voxels of the predicted and the ground truth segmentations. The values of the Dice scores are between 0 and 1, where a high value represents better prediction corresponding to the ground truth segmentation. We also assessed probabilistic tissue segmentation prediction performance for each tissue type, *T*, by computing the probabilistic similarity index (PSI) (Anbeek et al., 2005), as:

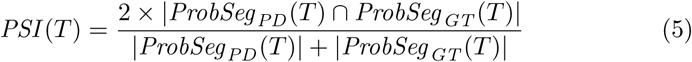

where |*ProbSeg* _*PD*_ *∩ ProbSeg* _*GT*_ |indicates the sum of the probability values over the intersection of the predicted and the ground truth segmentations, and |*ProbSeg* _*PD*_ (*T*)|and |*ProbSeg* _*GT*_ (*T*)|indicate the sum of the probability values of the predicted and the ground truth segmentations. Similar to the Dice score, the PSI is in the range between 0 and 1, where a higher value represents a better agreement with the ground truth. The mean and the standard deviation of the Dice score and PSI across the 10 testing HCP subjects and the 10 testing CAP datasets, respectively, were computed for a quantitative comparison.

## 4. Results

### 4.1. Comparison of tissue feature descriptors and machine learning classifiers

The comparison of *ACC* across the 9 combinations of 3 feature descriptors and 3 machine learning classifiers are presented in Tables 1 and 2 for the HCP and CAP data, respectively. The proposed classifier with augmented target loss function, *CNN* _*AT*_, obtained higher accuracy than both SVM and conventional CNN classifiers for each feature descriptor. The improvement in accuracy comparing to the conventional CNN was mostly in the tissue boundary region. Applying the proposed feature descriptor, *F*_*Proposed*_, which includes the 10 MK-Curve-based features, generated higher *ACC* than the other feature descriptors for each of the classifiers studied. Overall, applying our proposed classifier, *CNN* _*AT*_, on the proposed descriptor, *F*_*Proposed*_, achieved the highest tissue segmentation accuracy across the entire brain, with an *ACC* of 90.48% in the HCP data and 80.30% in the CAP data. Notably, the proposed *CNN* _*AT*_ also achieved reasonable accuracy using only the DTI features, *F*_*DTI*_ (Table 1 and Supplementary Fig. S1).

**Table 1:**
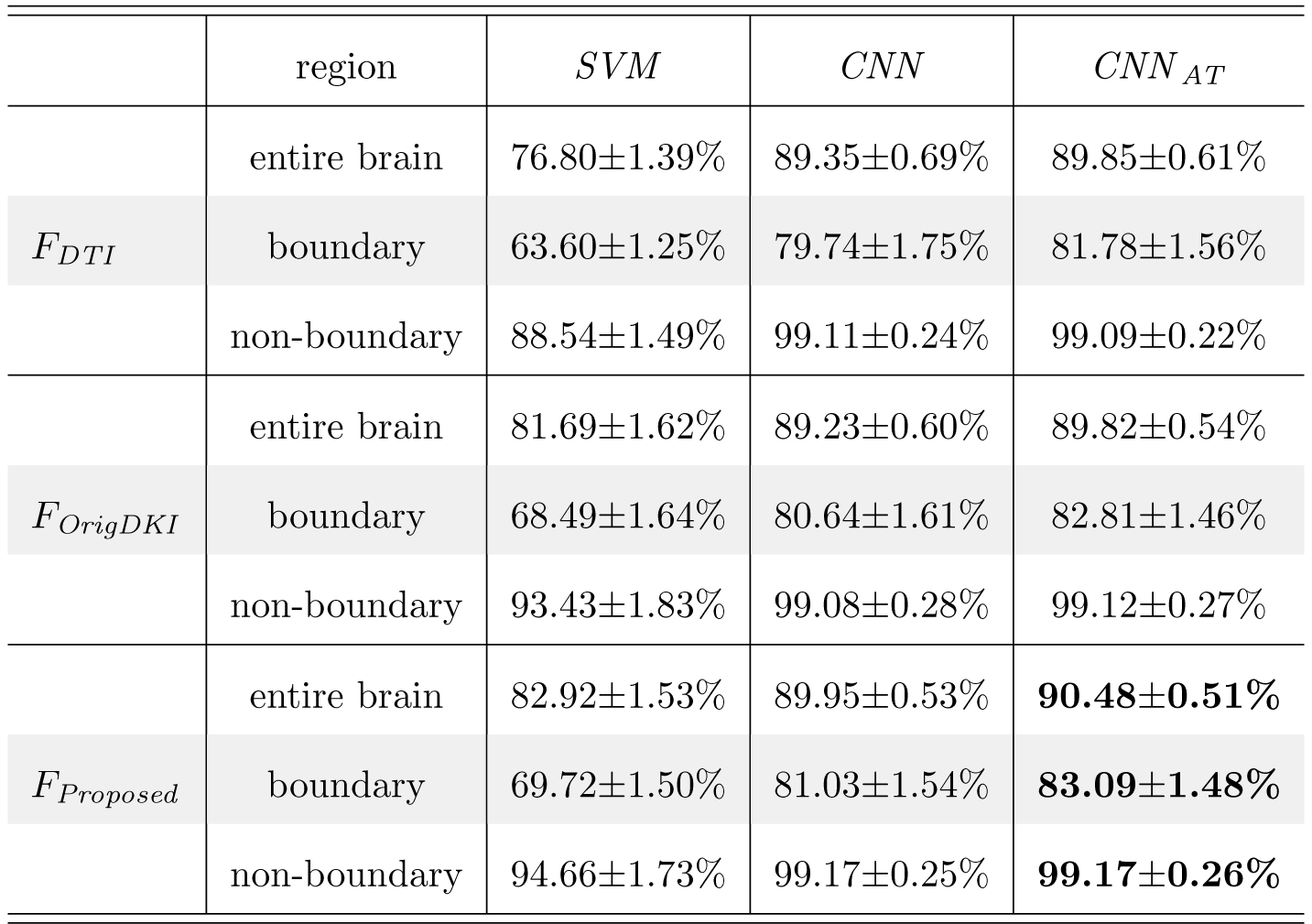
Quantitative comparison of the segmentation prediction accuracy across different tissue feature descriptors and machine learning classifiers on test HCP data. The prediction accuracy across the entire brain, and separately across tissue boundary and non-boundary regions are reported.

|  | region | *SVM* | *CNN* | *CNN<sub>AT</sub>* |
| --- | --- | --- | --- | --- |
|  | entire brain | 76.80±1.39% | 89.35±0.69% | 89.85±0.61% |
| *F<sub>DTI</sub>* | boundary | 63.60±1.25% | 79.74±1.75% | 81.78±1.56% |
|  | non-boundary | 88.54±1.49% | 99.11±0.24% | 99.09±0.22% |
|  | entire brain | 81.69±1.62% | 89.23±0.60% | 89.82±0.54% |
| *F<sub>OrigDKI</sub>* | boundary | 68.49±1.64% | 80.64±1.61% | 82.81±1.46% |
|  | non-boundary | 93.43±1.83% | 99.08±0.28% | 99.12±0.27% |
|  | entire brain | 82.92±1.53% | 89.95±0.53% | **90.48±0.51%** |
| *F<sub>Proposed</sub>* | boundary | 69.72±1.50% | 81.03±1.54% | **83.09±1.48%** |
|  | non-boundary | 94.66±1.73% | 99.17±0.25% | **99.17±0.26%** |

**Table 2:** Quantitative comparison of the segmentation prediction accuracy across different tissue feature descriptors and machine learning classifiers on the CAP data. The prediction accuracy across the entire brain, and separately across tissue boundary and non-boundary regions are reported.

|  | region | *SVM* | *CNN* | *CNN<sub>AT</sub>* |
| --- | --- | --- | --- | --- |
|  | entire brain | 56.65±6.61% | 76.25±3.66% | 76.68±4.35% |
| *F<sub>DTI</sub>* | boundary | 55.82±6.62% | 66.22±4.21% | 69.49±4.64% |
|  | non-boundary | 57.58±6.62% | 88.78±3.56% | 90.32±2.79% |
|  | entire brain | 59.55±3.53% | 77.78±3.10% | 78.35±3.02% |
| *F<sub>OrigDKI</sub>* | boundary | 58.73±3.54% | 66.19±3.67% | 70.36±3.67% |
|  | non-boundary | 60.48±3.52% | 91.53±2.14% | 91.57±2.30% |
|  | entire brain | 62.28±3.49% | 79.39±2.25% | **80.30±2.02%** |
| *F<sub>Proposed</sub>* | boundary | 61.46±3.48% | 68.70±3.05% | **73.00±2.78%** |
|  | non-boundary | 63.21±3.48% | 92.05±2.45% | **92.62±2.55%** |

### 4.2. Comparison with state-of-the-art tissue segmentation methods

Quantitative comparison (Fig. 2) shows that our method consistently out-performs the three state-of-the-art methods (CSD, SPM+b0 and daDWI), across the three evaluation measures (*ACC, Dice*, and *PSI*) and across the three tissue types (WM, GM and CSF).

**Figure 2:**
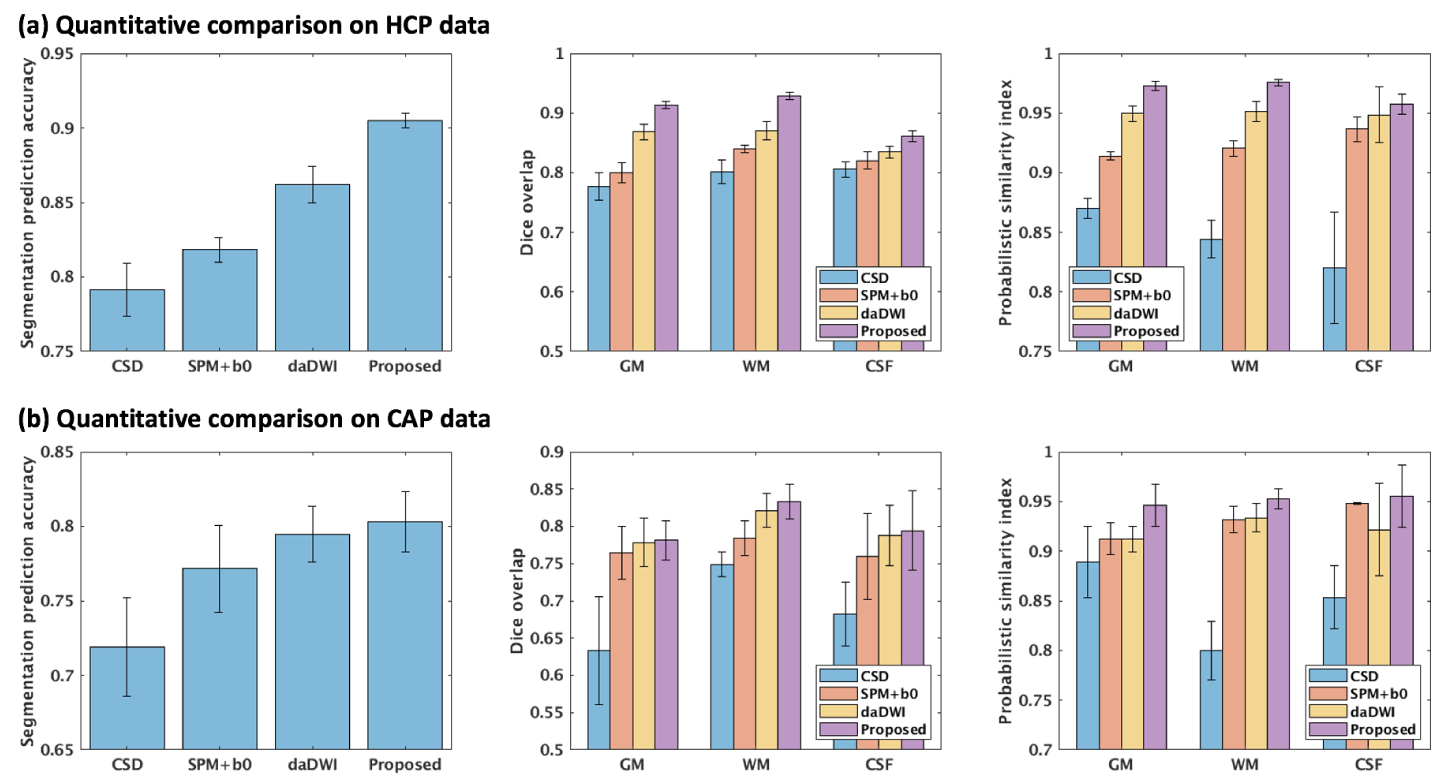
Quantitative comparison of the overall segmentation prediction accuracy, the Dice overlap, and the probabilistic similarity index (PSI) across the CSD, the SPM+b0, the daDWI, and the proposed tissue segmentation methods. The daDWI method only outputs a binary mask for the CSF, without providing a probabilistic prediction, thus the PSI for the CSF in this method is not applicable.

Visual assessment of the predicted tissue segmentation and probabilistic maps for the HCP (see an example case in Fig. 3) and CAP (see an example case in Fig. 4) datasets shows that our method generates tissue segmentation maps that are highly visually similar to the T2w-based tissue segmentation. The daDWI method also obtains visually similar results to the T2w-based method, while the SPM+b0 and CSD methods are relatively less similar. For example, in Fig. 3, the SPM-b0 and the CSD methods mislabeled parts of the caudate as WM, and in Fig. 4, the SPM-b0 method mispredicted the putamen to be WM.

**Figure 3:**
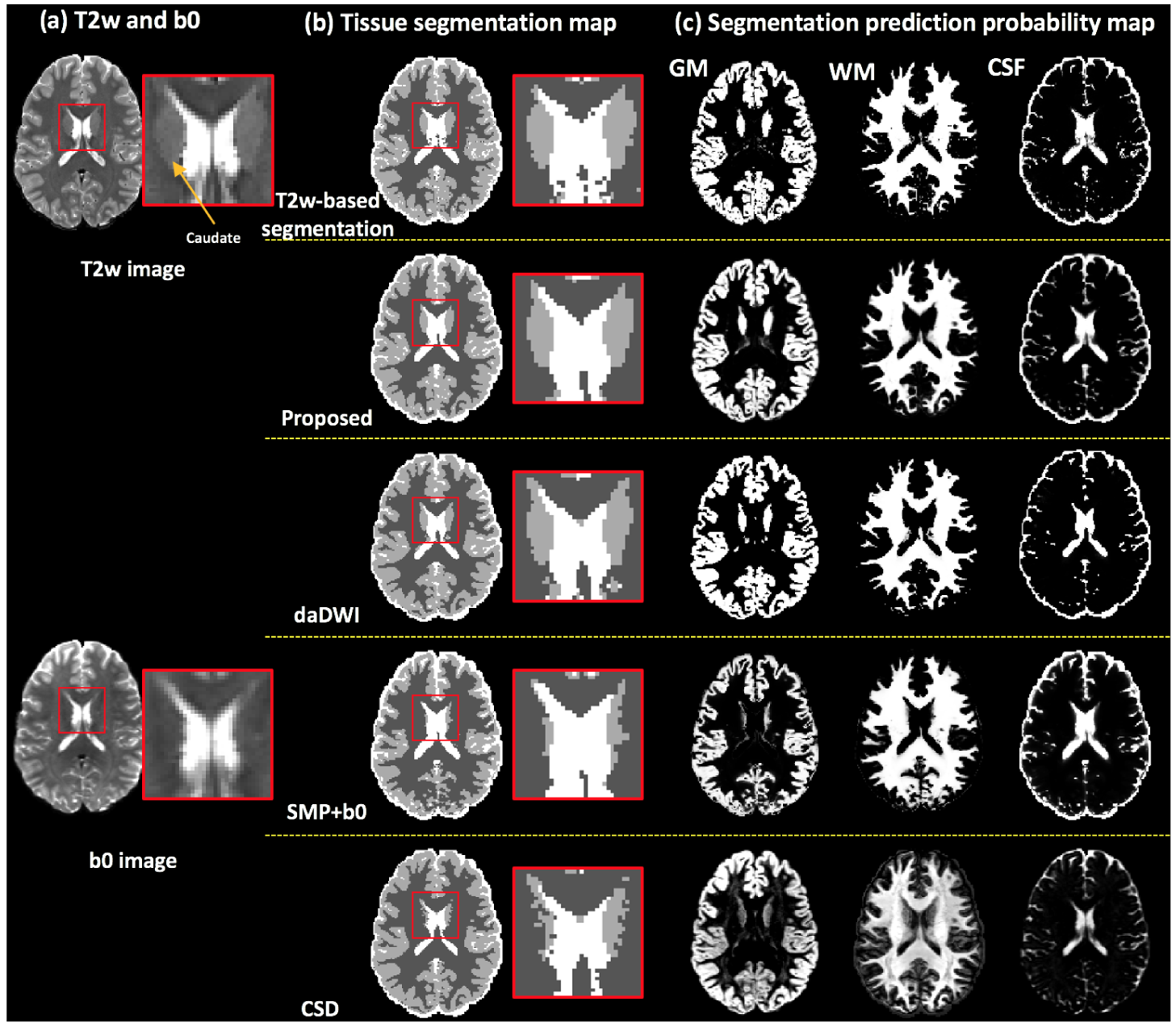
Visual comparison across the proposed method, the daDWI method, the SPM+b0 method and the CSD method on one example HCP subject, with respect to the T2w-based segmentation. (a) gives the T2w and the b0 images of the example HCP subject. (b) shows the comparison of the overall GM/WM/CSF segmentation. An inset image, enlarging part of the segmentation of the region near the caudate, is provided. (c) shows the comparison of the segmentation prediction probability map for each tissue type.

**Figure 4:**
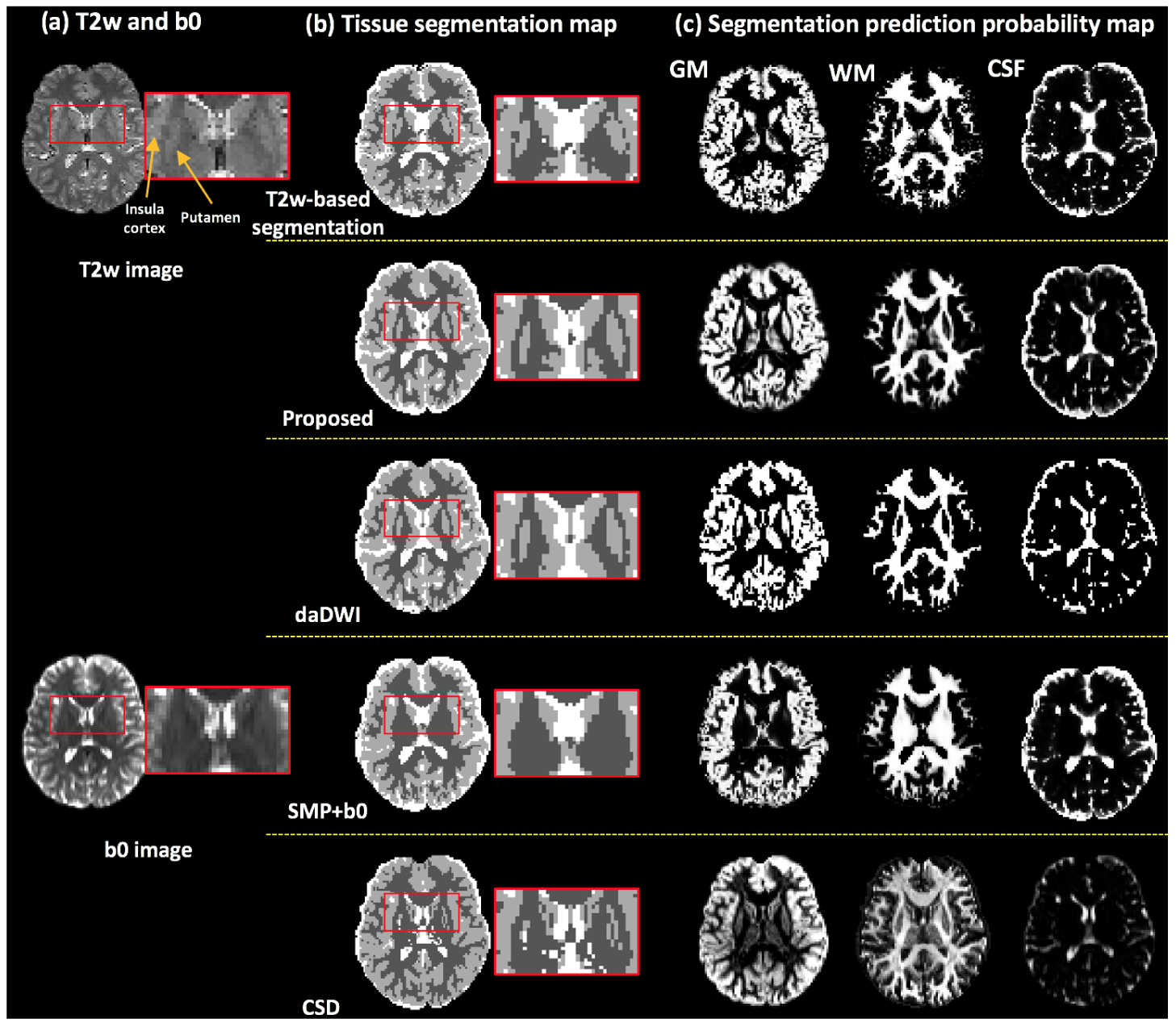
Visual comparison across the proposed method, the daDWI method, the SPM+b0 method and the CSD method on one example CAP subject, with respect to the T2w-based tissue segmentation. (a) gives the T2w and the b0 images of the example CAP subject. (b) shows the comparison of the overall GM/WM/CSF segmentation. An inset image, enlarging part of the segmentation of the region near the putamen, is provided. (c) shows the comparison of the segmentation prediction probability map for each tissue type.

Visual inspection also demonstrates that there are apparent tissue segmentation errors in the T2w-based method, in particular in the CAP data, whereas our method achieves a more anatomically correct tissue segmentation. For example, in the T2w-based tissue segmentation in Fig. 4(b), several WM voxels belonging to the external capsule between the putamen and the insula cortex are mislabeled to be GM. Our method generates a tissue segmentation map that is smoother and more closely resembles the expected shape of the putamen and the insula cortex, better corresponding with the anatomy as appearing on the T2w image.

## 5. Discussion

In this work, we proposed a novel deep learning brain tissue segmentation method that can be applied directly on dMRI data. We trained a CNN tissue segmentation model from high quality HCP data, using MK-Curve-based DKI features and a new augmented target loss function. On the testing HCP data, our method achieved highly comparable results to anatomical T2w-based “ground truth” tissue segmentation, while avoiding inter-modality registration. We also demonstrated that our method provided a good ability to generalize to dMRI data from a different acquisition with lower spatial and angular resolutions than those of the training HCP data. Quantitative and visual comparisons showed that our method outperformed several state-of-the-art dMRI-based tissue segmentation methods.

The two CNN-based classifiers (i.e., with the conventional categorical cross-entropy loss function and with the augmented target loss function) obtained much higher prediction accuracies than the SVM classifier. The improvement might be because CNN involves application of spatial operators that in each voxel take into account information from neighboring voxels, while the SVM classifier performs prediction based only on features derived from each individual voxel. Therefore, the CNN classifier can identify textures, shapes and relative locations of the voxels in the input feature volume that could contribute to the segmentation task. Comparing between the two CNN-based classifiers, incorporating the augmented target loss function generated a higher prediction accuracy, in particular on the tissue boundary region, which suggests that additional penalizations on misclassification of voxels at the tissue boundaries and the CSF tissue type was important for better segmentation.

The proposed *CNN* _*AT*_ achieved the highest prediction accuracy when using the 10 MK-Curve-based feature descriptor. However, the feature descriptor with only the DTI parameters also showed a large improvement compared with the SVM classifier with a promising prediction accuracy (89.85% on the HCP data and 76.68% on the CAP data; see Supplementary Fig. S2 for a visualization of DTI-feature-based tissue segmentation). These results demonstrate that the proposed CNN-based segmentation is also useful for standard single-shell dMRI data, where the additional DKI parameters are not available. However, when multi-shell data is available, adding DKI parameters (before or after MK-Curve correction) improves the prediction accuracy when compared to using only the DTI parameters. The improvements were potentially due to including information about restricted water diffusion properties that could be inferred from DKI but not DTI, and that better segments white matter from CSF or GM. Correcting implausible parameters and adding the MK-Curve features consistently obtained a higher prediction accuracy compared to the original DKI parameters, demonstrating the usefulness of the MK-Curve method for tissue segmentation.

We showed good performance of the proposed method when applied to the clinical acquisition data, despite the fact that this data had a lower image resolution and a lower number of gradient directions than the training HCP data. The ability of a tissue segmentation method to generalize to data from different acquisitions is important. dMRI acquisitions can have widely varying numbers of gradient directions, b-values, and magnitude of b-values, posing a challenge for machine learning. In our quantitative evaluation we showed that nearly 80% of the voxels were in agreement with the SPM segmentation of the coregistered T2w data, which was higher than the other methods that were evaluated.

Our proposed method performed tissue segmentation prediction directly from the dMRI data and thus could avoid obvious segmentation errors when transferring the anatomical T2w-based “ground truth” segmentation to the dMRI space. In the literature, anatomical-MRI-based segmentation, e.g., the one obtained by SPM, is usually used as the “ground truth” data (Ciritsis et al., 2018; Schnell et al., 2009; Cheng et al., 2020), since the segmentation appears in good agreement with the known anatomy. However, transferring T1w- or T2w-based segmentation into the dMRI space is challenging due to the image distortions in dMRI data, which affected inter-modality registration significantly (Albi et al., 2018; Wu et al., 2008; Jones and Cercignani, 2010). In addition, anatomical MRI data often has higher spatial resolution than dMRI data, resulting in segmentation errors from smoothing and image interpolation when transferring the tissue segmentation computed from the high-resolution anatomical data to the low-resolution dMRI data. Segmentation errors in the “ground truth” T2w-based segmentation were also reflected in our data, especially in the CAP data where visual inspection of the T2w co-registered segmentation appears to mislabel some brain areas, in particular at the tissue boundaries. We showed that our proposed method was able to correctly segment in areas where the T2w coregistered segmentation were apparently wrong, thus generating a visually more correct segmentation corresponding to the anatomy as appearing on the T2w image. Similar to previous studies (Beejesh et al., 2019; Hui et al., 2015; Steven et al., 2014; Cheng et al., 2020; Yap et al., 2015), we chose to calculate the accuracy against the “ground truth” segmentation from co-registered anatomical MRI data. Therefore, regions where our method correctly labeled the tissue type but the “ground truth” did not would cause a lower accuracy score. This observation might explain the overall relatively lower accuracy scores of the CAP data, where acquisition with reversed phase encoding was not available and image resolution was relatively low compared to the HCP data.

Potential future directions and limitations of the current work are as follows. First, the current work focused on tissue segmentation of the GM, WM and CSF. An interesting future investigation could include segmentation of more specific anatomical structures such as subcortical GM regions, white matter bundles, and specific cortices. Second, in the current work, we evaluated our method on healthy adult brains. Future work could include investigation of the proposed tissue segmentation method in brains with lesions (such as tumors or edema) and/or from different age ranges (e.g. from children and elderly people). Such work would require curation of further training data that reflect anatomy of certain populations. Third, in the present study, we explored a Unet architecture, which provided highly promising performance. Future work could include an investigation of more advanced network architectures, in combination with our proposed augmented target loss function to further improve classification accuracy.

## Acknowledgment

The authors would like to thank the NYU Diffusion MRI group (DKI model fit), the SPM community, the MRtrix community (CSD tissue segmentation) for making code used in this study available online, and Dr. Hu Cheng for sharing the daDWI segment code. We gratefully acknowledge funding provided by the following National Institutes of Health (NIH) grants: P41 EB015902, P41 EB015898, R01 MH108574, R01 MH074794, R01 MH119222, R21 MH115280, and U01 CA199459.

## Footnotes

1 In the CSD framework, a response function is used as the kernel during the deconvolution step to estimate the fiber orientation distribution (FOD) function.

## References

Akkus, Z., Galimzianova, A., Hoogi, A., Rubin, D.L., Erickson, B.J., 2017. Deep learning for brain MRI segmentation: state of the art and future directions. Journal of digital imaging 30, 449–459.

Albi, A., Meola, A., Zhang, F., Kahali, P., Rigolo, L., Tax, C.M., Ciris, P.A., Essayed, W.I., Unadkat, P., Norton, I., Rathi, Y., Olubiyi, O., Golby, A.J., O’Donnell, L.J., 2018. Image Registration to Compensate for EPI Distortion in Patients with Brain Tumors: An Evaluation of Tract-Specific Effects. Journal of Neuroimaging 28, 173–182.

Anbeek, P., Vincken, K.L., Van Bochove, G.S., Van Osch, M.J., van der Grond, J., 2005. Probabilistic segmentation of brain tissue in MR imaging. Neuroimage 27, 795–804.

Ashburner, J., Friston, K.J., 2005. Unified segmentation. Neuroimage 26, 839–851.

Avants, B.B., Tustison, N., Song, G., 2009. Advanced normalization tools (ANTS). Insight j 2, 1–35.

Beejesh, A., Gopi, V.P., Hemanth, J., 2019. Brain MR kurtosis imaging study: Contrasting gray and white matter. Cognitive Systems Research 55, 135–145.

Bernal, J., Kushibar, K., Asfaw, D.S., Valverde, S., Oliver, A., Martí, R., Lladó, X., 2019. Deep convolutional neural networks for brain image analysis on magnetic resonance imaging: a review. Artificial intelligence in medicine 95, 64–81.

Breger, A., Orlando, J., Harar, P., Dörfler, M., Klimscha, S., Grechenig, C., Gerendas, B., Schmidt-Erfurth, U., Ehler, M., 2020. On Orthogonal Projections for Dimension Reduction and Applications in Augmented Target Loss Functions for Learning Problems. Journal of Mathematical Imaging and Vision 62, 376 – 394.

Cheng, H., Newman, S., Afzali, M., Fadnavis, S.S., Garyfallidis, E., 2020. Segmentation of the brain using direction-averaged signal of DWI images. Magnetic Resonance Imaging 69, 1–7.

Ciritsis, A., Boss, A., Rossi, C., 2018. Automated pixel-wise brain tissue segmentation of diffusion-weighted images via machine learning. NMR in Biomedicine 31, e3931.

Dhollander, T., Raffelt, D., Connelly, A., 2016. Unsupervised 3-tissue response function estimation from single-shell or multi-shell diffusion mr data without a co-registered t1 image, in: ISMRM, p. 5.

Dhollander, T., Raffelt, D., Connelly, A., 2018. Accuracy of response function estimation algorithms for 3-tissue spherical deconvolution of diverse quality diffusion MRI data, in: ISMRM, p. 1569.

Dong, H., Yang, G., Liu, F., Mo, Y., Guo, Y., 2017. Automatic brain tumor detection and segmentation using u-net based fully convolutional networks, in: Annual Conference on Medical Image Understanding and Analysis, Springer. pp. 506–517.

Fischl, B., 2012. FreeSurfer. NeuroImage 62, 774–781.

Garcia-Garcia, A., Orts-Escolano, S., Oprea, S., Villena-Martinez, V., Garcia-Rodriguez, J., 2017. A review on deep learning techniques applied to semantic segmentation. 1704.06857.

Glasser, M.F., Sotiropoulos, S.N., Wilson, J.A., Coalson, T.S., Fischl, B., Andersson, J.L., Xu, J., Jbabdi, S., Webster, M., Polimeni, J.R., et al., 2013. The minimal preprocessing pipelines for the Human Connectome Project. Neuroimage 80, 105–124.

Goodfellow, I., Bengio, Y., Courville, A., 2016. Deep learning. MIT press.

Hui, E.S., Glenn, G.R., Helpern, J.A., Jensen, J.H., 2015. Kurtosis analysis of neural diffusion organization. Neuroimage 106, 391–403.

Hwang, H., Rehman, H.Z.U., Lee, S., 2019. 3D U-Net for skull stripping in brain MRI. Applied Sciences 9, 569.

Jenkinson, M., Beckmann, C.F., Behrens, T.E., Woolrich, M.W., Smith, S.M., 2012. Fsl. Neuroimage 62, 782–790.

Jensen, J.H., Helpern, J.A., 2010. MRI quantification of non-Gaussian water diffusion by kurtosis analysis. NMR in Biomedicine 23, 698–710.

Jeurissen, B., Tournier, J.D., Dhollander, T., Connelly, A., Sijbers, J., 2014. Multi-tissue constrained spherical deconvolution for improved analysis of multi-shell diffusion MRI data. NeuroImage 103, 411–426.

Jones, D.K., Cercignani, M., 2010. Twenty-five pitfalls in the analysis of diffusion MRI data. NMR in Biomedicine 23, 803–820.

Krizhevsky, A., Sutskever, I., Hinton, G.E., 2012. Imagenet classification with deep convolutional neural networks, in: Advances in neural information processing systems, pp. 1097–1105.

Kumazawa, S., Yoshiura, T., Honda, H., Toyofuku, F., 2013. Improvement of partial volume segmentation for brain tissue on diffusion tensor images using multiple-tensor estimation. Journal of digital imaging 26, 1131–1140.

Kumazawa, S., Yoshiura, T., Honda, H., Toyofuku, F., Higashida, Y., 2010. Partial volume estimation and segmentation of brain tissue based on diffusion tensor MRI. Medical physics 37, 1482–1490.

LeCun, Y., Bottou, L., Bengio, Y., Haffner, P., 1998. Gradient-based learning applied to document recognition. Proceedings of the IEEE 86, 2278–2324.

Liu, T., Li, H., Wong, K., Tarokh, A., Guo, L., Wong, S.T., 2007. Brain tissue segmentation based on DTI data. NeuroImage 38, 114–123.

Mah, Y.H., Jager, R., Kennard, C., Husain, M., Nachev, P., 2014. A new method for automated high-dimensional lesion segmentation evaluated in vascular injury and applied to the human occipital lobe. Cortex 56, 51–63.

Malinsky, M., Peter, R., Hodneland, E., Lundervold, A., Lundervold, A., Jan, J., 2013. Registration of FA and T1-weighted MRI data of healthy human brain based on template matching and normalized cross-correlation. J Digit Imaging 26, 774–785.

McKinnon, E.T., Jensen, J.H., Glenn, G.R., Helpern, J.A., 2017. Dependence on b-value of the direction-averaged diffusion-weighted imaging signal in brain. Magnetic resonance imaging 36, 121–127.

Nie, D., Wang, L., Adeli, E., Lao, C., Lin, W., Shen, D., 2018. 3-D fully convolutional networks for multimodal isointense infant brain image segmentation. IEEE transactions on cybernetics 49, 1123–1136.

Norton, I., Essayed, W.I., Zhang, F., Pujol, S., Yarmarkovich, A., Golby, A.J., Kindlmann, G., Wassermann, D., Estepar, R.S.J., Rathi, Y., et al., 2017. SlicerDMRI: open source diffusion MRI software for brain cancer research. Cancer research 77, e101–e103.

O’Donnell, L.J., Pasternak, O., 2015. Does diffusion MRI tell us anything about the white matter? An overview of methods and pitfalls. Schizophrenia research 161, 133–141.

Parker, J.R., 2010. Algorithms for image processing and computer vision. John Wiley & Sons.

Ronneberger, O., Fischer, P., Brox, T., 2015. U-net: Convolutional networks for biomedical image segmentation, in: International Conference on Medical image computing and computer-assisted intervention, Springer. pp. 234–241.

Schnell, S., Saur, D., Kreher, B., Hennig, J., Burkhardt, H., Kiselev, V., 2009. Fully automated classification of HARDI in vivo data using a support vector machine. NeuroImage 46, 642 – 651.

Shaw, C.B., Jensen, J.H., 2017. Recent Computational Advances in Denoising for Magnetic Resonance Diffusional Kurtosis Imaging (DKI). Journal of the Indian Institute of Science 97, 377–390.

Smith, S.M., Jenkinson, M., Woolrich, M.W., Beckmann, C.F., Behrens, T.E., Johansen-Berg, H., Bannister, P.R., Luca, M.D., Drobnjak, I., Flitney, D.E., Niazy, R.K., Saunders, J., Vickers, J., Zhang, Y., Stefano], N.D., Brady, J.M., Matthews, P.M., 2004. Advances in functional and structural MR image analysis and implementation as FSL. NeuroImage 23, S208 – S219.

Steven, A.J., Zhuo, J., Melhem, E.R., 2014. Diffusion kurtosis imaging: an emerging technique for evaluating the microstructural environment of the brain. American journal of roentgenology 202, W26–W33.

Tabesh, A., Jensen, J.H., Ardekani, B.A., Helpern, J.A., 2011. Estimation of tensors and tensor-derived measures in diffusional kurtosis imaging. Magnetic resonance in medicine 65, 823–836.

Tournier, J.D., Calamante, F., Connelly, A., 2007. Robust determination of the fibre orientation distribution in diffusion MRI: non-negativity constrained super-resolved spherical deconvolution. Neuroimage 35, 1459–1472.

Veraart, J., Fieremans, E., Jelescu, I.O., Knoll, F., Novikov, D.S., 2016. Gibbs ringing in diffusion MRI. Magnetic resonance in medicine 76, 301–314.

Wasserthal, J., Neher, P., Maier-Hein, K.H., 2018. Tractseg-fast and accurate white matter tract segmentation. NeuroImage 183, 239–253.

Wen, Y., He, L., von Deneen, K.M., Lu, Y., 2013. Brain tissue classification based on DTI using an improved Fuzzy C-means algorithm with spatial constraints. Magnetic Resonance Imaging 31, 1623 – 1630.

Wu, M., Chang, L.C., Walker, L., Lemaitre, H., Barnett, A.S., Marenco, S., Pierpaoli, C., 2008. Comparison of EPI distortion correction methods in diffusion tensor mri using a novel framework, in: Medical Image Computing and Computer-Assisted Intervention, pp. 321–329.

Yap, P.T., Zhang, Y., Shen, D., 2015. Brain Tissue Segmentation Based on Diffusion MRI Using L0 Sparse-Group Representation Classification, in: Medical Image Computing and Computer-Assisted Intervention, pp. 132–139.

Zhang, F., Ning, L., O’Donnell, L.J., Pasternak, O., 2019. MK-curve - Characterizing the relation between mean kurtosis and alterations in the diffusion MRI signal. NeuroImage 196, 68 – 80.

Zhang, F., Noh, T., Juvekar, P., Frisken, S.F., Rigolo, L., Norton, I., Kapur, T., Pujol, S., Wells, W., Yarmarkovich, A., Kindlmann, G., Wassermann, D., San Jose Estepar, R., Rathi, Y., Kikinis, R., Johnson, H.J., Westin, C.F., Pieper, S., Golby, A.J., O’Donnell, L.J., 2020. SlicerDMRI: Diffusion MRI and Tractography Research Software for Brain Cancer Surgery Planning and Visualization. JCO Clinical Cancer Informatics 4, 299–309.

Zhang, W., Li, R., Deng, H., Wang, L., Lin, W., Ji, S., Shen, D., 2015. Deep convolutional neural networks for multi-modality isointense infant brain image segmentation. NeuroImage 108, 214 – 224.

