## supplementary material for "Deep Learning Based Segmentation of Brain Tissue from Diffusion MRI"

### Supplementary materials

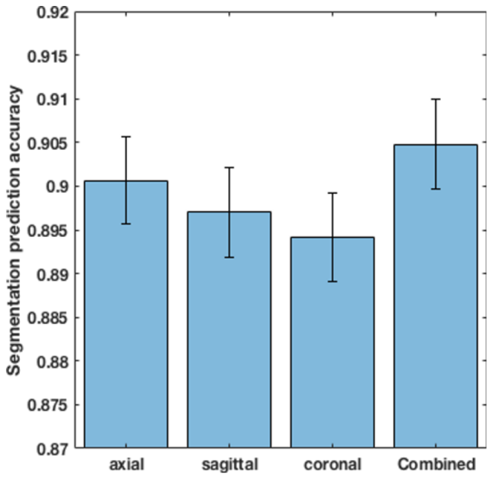


Fig. 1. Comparison of tissue segmentation prediction accuracy (ACC) using the Unet models trained from an individual view (the axial, sagittal, or coronal views) and the combination of all three views. Here, the mean accuracy and standard deviation of the ACC from the 10 HCP testing subjects are reported.

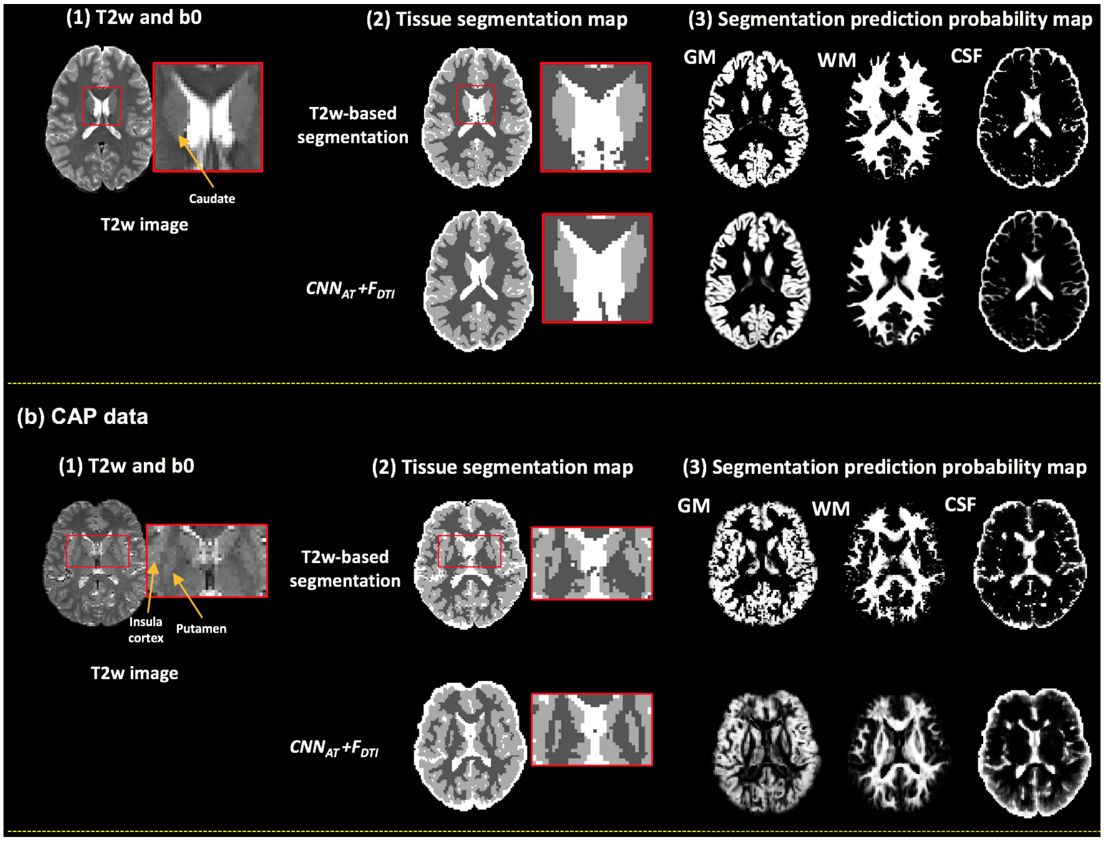


Fig. 2. Tissue segmentation prediction using the proposed $CNN_{AT}$ classifier, in combination with the $F_{DTI}$ feature descriptor on one example HCP dataset (a) and one example CAP dataset (b). Subfigures (a1, b1) show the T2w image of the example subjects. Subfigures (a2, b2) show the overall GM/WM/CSF segmentation, with comparison to the T2w-based segmentation. The inset shows an enlargment of the segmentation of the region near the caudate Subfigures (a3, b3) show the segmentation prediction probability map for each tissue type. Overall, using the proposed $CNN_{AT}$ classifier and DTI features generated comparable tissue segmentation to the T2w-based method.
